# Impact of central carbon metabolism bypasses on the production of beta-carotene in *Yarrowia lipolytica*

**DOI:** 10.1101/2023.11.10.566616

**Authors:** Tadej Markuš, Mladen Soldat, Vasilka Magdevska, Jaka Horvat, Martin Kavšček, Gregor Kosec, Štefan Fujs, Uroš Petrovič

## Abstract

*Yarrowia lipolytica* is an oleaginous yeast with ever growing popularity in the metabolic engineering circles. It is well known for its ability to accommodate a high carbon flux through acetyl-CoA and is being extensively studied for production of chemicals derived from it.

We investigated the effects of modifying the upstream metabolism leading to acetyl-CoA on beta-carotene production, including its titer, yield, and content. We examined the pyruvate and the phosphoketolase bypass, both of which are stoichiometrically favorable for the production of acetyl-CoA and beta-carotene. Additionally, we examined a set of genes involved in the carnitine shuttle. We constructed a set of parental strains derived from the *Y. lipolytica* YB-392 wild-type strain, each with a different capacity for beta-carotene production, and introduced genes for the metabolic bypasses in each of the constructed parental strains. Subsequently, we subjected these constructed strains to a series of fermentation experiments.

We discovered that altering the upstream metabolism in most cases led to a decrease in performance for production of beta-carotene. Most notably, a set of genes used for the pyruvate bypass (*YlPDC2*, *YlALD5*, and *YlACS1*) and the phosphoketolase bypass (*LmXPK* and *CkPTA*) resulted in the reduction of more than 30%.

Our findings contribute to our understanding of *Y. lipolytica*’s metabolic capacity and suggest that production of beta-carotene is most likely not limited solely by the acetyl-CoA supply. We also highlight a complex nature of engineering *Y. lipolytica*, as most of the results from studies using a different strain background did not align with our findings.

## Introduction

*Yarrowia lipolytica* is an oleaginous yeast known for its remarkable lipid accumulation capacity, with some engineered strains reportedly reaching lipid content of 90% of dry weight and above (1, 2). It exhibits versatile substrate utilization, with recent advancements in metabolic engineering allowing it to utilize lignocellulosic materials like xylulose (3). *Y. lipolytica* demonstrates exceptional growth potential in high-density industrial fermenter environments (4), rendering it highly suitable for various industrial applications. Moreover, the development of genetic tools specific to *Y. lipolytica* has experienced significant progress in recent years (5), although not reaching the same level of advancement as its relatively closely related counterpart, *Saccharomyces cerevisiae*. The status of *Y. lipolytica* as an industrial workhorse is further reinforced by its Generally Recognized as Safe (GRAS) designation by the American Food and Drug Administration (FDA) (GRN No. 355, 632, 759, 940) and Qualified Presumption of Safety (QPS) status by the European Food Safety Authority (EFSA) (6).

Most of the studies on *Y. lipolytica* in the field of biotechnology mainly focus on products derived from acetyl coenzyme A (acetyl-CoA), a key molecule in the production of lipids and lipophilic commodities. Acetyl-CoA serves as a common precursor to terpenoids, which represent the largest and the most diverse class of natural molecules (7) and have garnered increasing attention from the industry. In order to enhance the production of a desired bioproduct, it is essential to direct a significant amount of carbon flux towards its biosynthesis. This can be achieved through two main strategies: “pulling” carbon towards the end product, or “pushing” carbon through increased substrate supply. In our study, we focused on evaluating different “push” strategies to drive carbon towards our desired end-product – carotenoids. Carotenoids serve as precursors to various commercially valuable natural compounds, including vitamin A, saffron compounds, and phytohormones. The development of a commercially viable bioprocess for carotenoid production necessitates achieving specific key performance indicators (KPIs), typically encompassing titer, rate, and yield. In this study, our objective was to evaluate the impact of different metabolic bypasses on the KPIs for the biotechnological production of beta-carotene.

To assess the impact of increased substrate supply, we examined three metabolic pathways that lead to the production of acetyl-CoA in *Y. lipolytica*: (i) the pyruvate bypass, (ii) the phosphoketolase bypass, and (iii) the carnitine shuttle system (Fig 1). It is noteworthy that *Y. lipolytica* already possesses a natural ability for high production of acetyl-CoA (8, 9). By overexpressing the genes for these three pathways alongside *Y. lipolytica*’s natural ability for acetyl-CoA production, we sought to evaluate the effectiveness of enhancing substrate supply on the production of beta-carotene.

**Fig 1:**
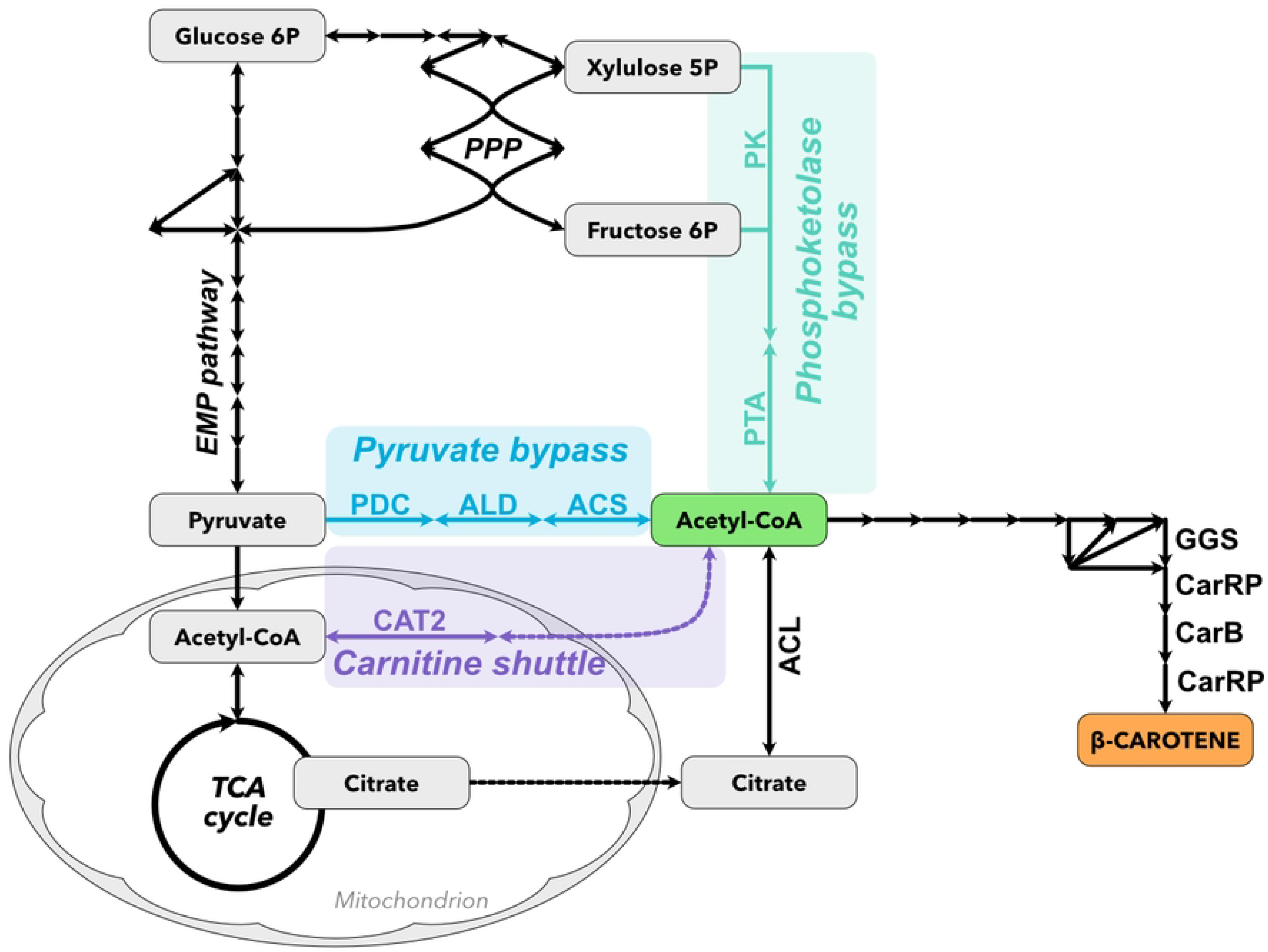
Schematic representation of the central carbon metabolism and modifications leading to cytosolic acetyl-CoA. ACL: ATP citrate lyase, ACS: acetyl-CoA synthetase, ALD: acetaldehyde dehydrogenase, CarB: phytoene desaturase, CarRP: bifunctional lycopene cyclase/phytoene synthase, CAT2: carnitine-O-acetyltransferase, GGS: geranylgeranyl pyrophosphate synthase, PDC: pyruvate decarboxylase, PK: phosphoketolase, PPP: pentose phosphate pathway, PTA: Phosphate acetyltransferase, TCA: tricarboxylic acid cycle.

## Results and discussion

### Selecting strains and genes for metabolic bypasses

*Y. lipolytica* is renowned for its capacity to accumulate lipids as an energy storage mechanism. It can either assimilate lipids from the environment or synthesize triglycerides (TAGs) efficiently when ample carbon sources are available. However, there exists considerable variability in the lipid accumulation and synthesis abilities among individual *Y. lipolytica* strains (10). Although strain W29 is commonly used by numerous research groups, we opted to work with the strain YB-392 due to its superior lipid production capabilities (11). Lipid droplets serve as storage sites ideal for other lipophilic compounds, such as carotenoids. Studies have demonstrated a positive correlation between increased general lipid production and enhanced beta-carotene production (12). We prepared a set of strains to comprehensively investigate the impact of metabolic bypasses in different genetic backgrounds (Fig 2). These strains enabled us to compare the impacts of the introduced bypasses at different stages of strain engineering. By introducing two cassettes for the biosynthesis of beta-carotene, we effectively reached the upper limit of the cell’s capacity to produce beta-carotene using the “pull” strategy, since the introduction of a third cassette did not result in significantly increased beta-carotene production, suggesting that further enhancements may be limited by other factors (Fig 3).

**Fig 2:**
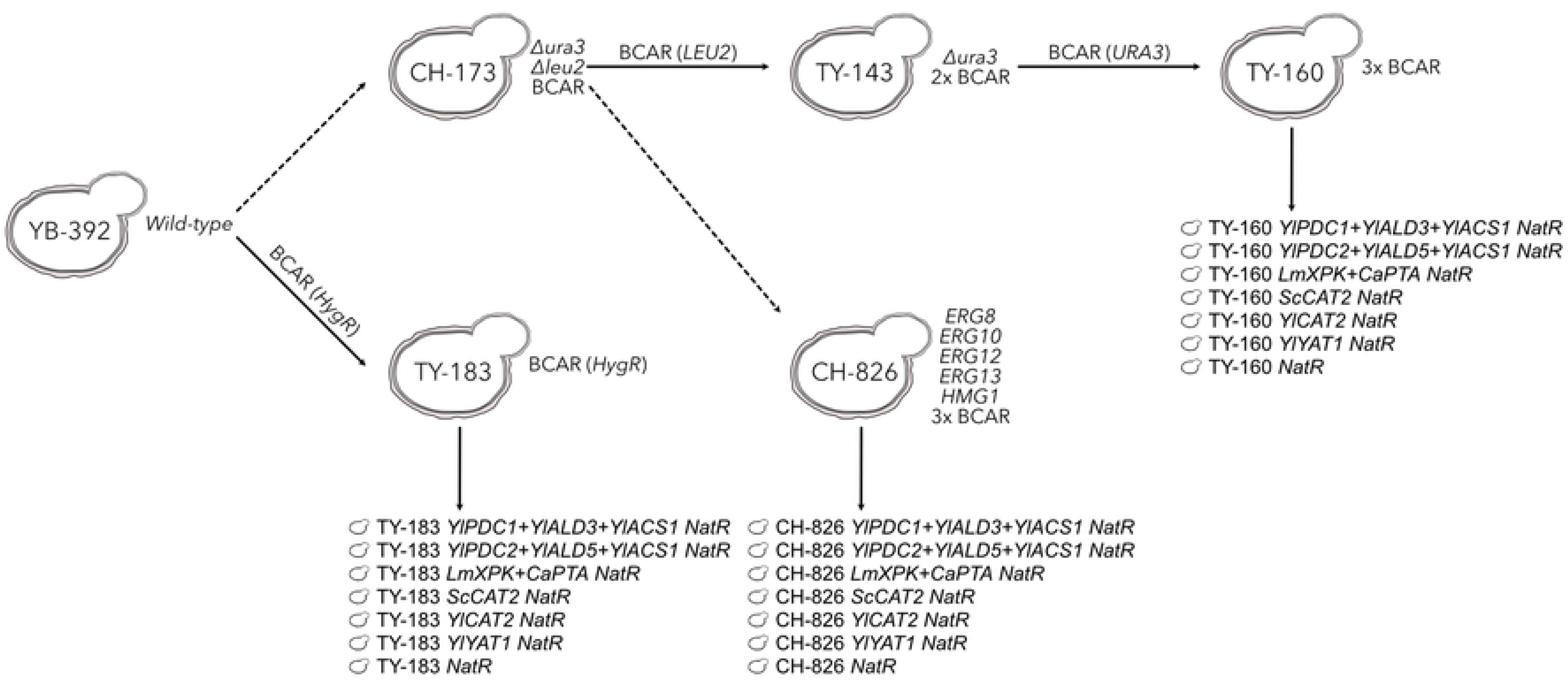
The lineage of parent strains used in the study. Dashed arrows represent multiple steps involved in the construction of the strain, while solid arrows indicate a single step. The YB-392 strain is a pure wild-type strain. The CH-173 strain is a derivative of YB-392, engineered for easier manipulation through deletion of *URA3* and *LEU2* genes. Additionally, it carries an overexpression cassette for the biosynthesis of beta-carotene (BCAR). The BCAR cassette comprises codon-optimized genes from *Mucor circinelloides* (*carB* and *carRP*) along with *Y. lipolytica*’s native gene *GGS1*. All genes are driven using the strong constitutive P_hp8d_ promoter and terminated with the T_cyc1_ terminator. The TY-160 strain was constructed by introducing two additional B-CAR cassettes, each designed to complement one auxotrophic marker. The TY-183 strain was created by transforming the YB-392 with a single BCAR cassette and using hygromycin as a selection marker. Finally, the CH-826 strain is a beta-carotene-producing strain with multiple overexpressed genes and mayor bottlenecks removed. We focused mainly on the prototrophic strains because we observed a considerate impact of auxotrophies on the production of beta-carotene (**Error! Reference source not found.**).

**Fig 3:**
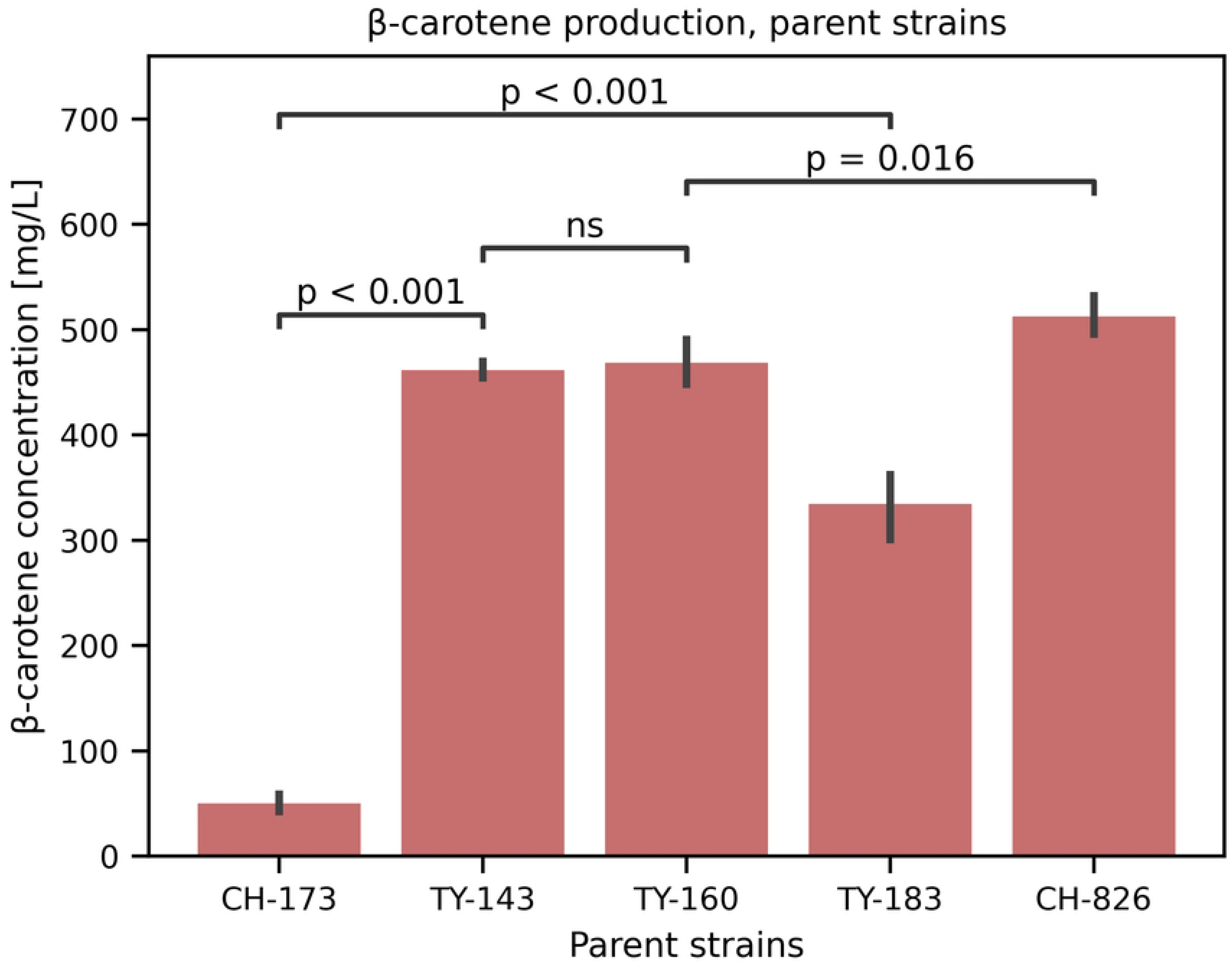
Beta-carotene production by the parental strains. Fermentation was performed in a 48-well plate in the SC-5 mineral medium. Error bars represent confidence interval of 95%. N=6 for each parent strains. Introducing a third overexpression cassette for beta-carotene biosynthesis does not enhance its production (TY-143 vs TY-160).

Our aim was to assess the contribution of modifications in the metabolic pathways that lead to the production of acetyl-CoA to increased carotenoid biosynthesis. We analyzed the contribution of complementary pathways for acetyl-CoA supply, in addition to the naturally occurring ATP citrate lyase (ACL) pathway, to enhance beta-carotene production. Increasing the substrate supply in the form of acetyl-CoA should result in improved lipid content and increased carotenoid production (12).

We selected three metabolic bypasses within the central carbon metabolism that lead to acetyl-CoA: (i) the pyruvate bypass, (ii) the phosphoketolase bypass, and (iii) the carnitine shuttle (Fig 1). The pyruvate and phosphoketolase bypasses are stoichiometrically favorable for acetyl-CoA and subsequent beta-carotene production, while the carnitine shuttle facilitates the transport of acetyl-CoA from the mitochondria. This way of acetyl-CoA export is hypothesized to be a response to metabolic flux overflow, thus preventing enzyme and protein damage via spontaneous acetylation in mitochondria (13). Some previous studies have demonstrated the benefits of such engineering approaches for producing compounds similar to beta-carotene (Table 1).

**Table 1:** Previous studies utilizing metabolic bypasses in *Y. lipolytica* for metabolic engineering.

| Approach | Genes | Product | Production | Reference |
| --- | --- | --- | --- | --- |
| Carnitine shuttle | *ScCAT2* | Lipids | 66.4 g/L | (14) |
| Phosphoketolase bypass | *LmXPK, CaPTA* | Fatty acid methyl esters | 98.9 g/L | (15) |
| Phosphoketolase bypass | *BbxfpK, AcxpkA* | Various aromatic chemical | 593.53 ± 28.75 mg/L (p-Coumaric acid) | (16) |
| Phosphoketolase bypass | *BbPK, BsPTA* | Beta-ionone | 358.4 mg/L | (17) |
| Phosphoketolase bypass | *TsPTA, CaXPK* | Lipids | ~5 g/L | (18) |
| Carnitine shuttle, phosphoketolase bypass | *ScCAT2, LmPK, CkPTA* | Alpha-humulene | 1413.2 ± 42.8 mg/L | (19) |
| Carnitine shuttle, phosphoketolase bypass | *ScCAT2, AnPK, BsuPTA* | Beta-farnesene | 28.9 g/L | (20) |
| Carnitine shuttle, phosphoketolase bypass, pyruvate bypass | *YICAT2, AnPK, BsuPTA, EcPDC, EcALDH, ScACS2* | Trans-nerolidol | 11.1 g/L | (21) |
| Pyruvate bypass | *YIPDC1, YIPDC2, YIALD1, YIALD2, YIALD3, YIALD4, YIALD5, YIALD6, YIACS1* | Triacetic lactone | 35.9 ± 3.85 g/L (bioreactor) | (22) |
| Pyruvate bypass | *YIPDC1, YIALD4* | Triacetic lactone | 4.76 g/L (Flask) | (23) |
| Pyruvate bypass | *YIACS, YIPDC1, YIALD2* | Squalene | 10 mg/g DCW | (24) |

### Combinatorial screening

At the initial stage of our study, we employed a combinatorial approach to explore various combinations of metabolic genes involved in the pyruvate bypass and the phosphoketolase bypass (Table 2). The objective was to identify the optimal gene combination for these two bypasses. For the combinatorial assembly we utilized an adapted YTK system (25). The YTK system enabled us to design the assembly in such a way that only plasmids containing one of each required genes could successfully assemble. The prepared mixture was then transformed into the CH-173 strain. We screened over 3500 individual transformed colonies and then selected some of the top performing strains for further evaluation. However, upon re-testing the strains, the results revealed no significant improvement over the control strains (Fig 4): on the contrary, most of the re-evaluated strains carrying either of the bypass pathways exhibited decreased beta-carotene production compared to the control. As a result, we shifted our focus towards a more in-depth investigation, narrowing down our analysis to a smaller subset of central carbon metabolism modifications, which are listed in Table 3.

**Table 2:** Metabolic genes involved in central carbon metabolism used in this study.

| Gene | Source organism | Enzyme function | Pathway | Accession No. | Reference |
| --- | --- | --- | --- | --- | --- |
| *ApPK* | *Aureobasidium pullulans* | Phosphoketolase | Phosphoketolase bypass | THX32035 | This study |
| *BalXFP* | *Bifidobacterium animalis* | Phosphoketolase | Phosphoketolase bypass | WP_012754421 | This study |
| *LmXFP* | *Leuconostoc mesenteroides* | Phosphoketolase | Phosphoketolase bypass | WP_002815143 | This study |
| *LmXPKA* | *Leuconostoc*<br>*mesenteroides* | Phosphoketolase | Phosphoketolase<br>bypass | WP_011680497 | (15, 17, 19) |
| *LpXPKA* | *Lactiplantibacillus*<br>*pentosus* | Phosphoketolase | Phosphoketolase<br>bypass | WP_050339622 | This study |
| *SpPK* | *Schizosaccharomyces*<br>*pombe* | Phosphoketolase | Phosphoketolase<br>bypass | NP_595963 | This study |
| *TIPK* | *Trichoderma*<br>*longibrachiatum* | Phosphoketolase | Phosphoketolase<br>bypass | PTB75968 | This study |
| *CkPTA* | *Clostridium kluyveri* | Phosphate<br>acetyltransferase | Phosphoketolase<br>bypass | WP_012101779 | (15, 17, 19) |
| *DrPTA* | *Deinococcus*<br>*radiodurans* | Phosphate<br>acetyltransferase | Phosphoketolase<br>bypass | AAF09663 | This study |
| *EcPTA* | *Escherichia coli* | Phosphate<br>acetyltransferase | Phosphoketolase<br>bypass | BAA04502 | This study |
| *PaPTA* | *Pseudoalteromonas sp.* | Phosphate<br>acetyltransferase | Phosphoketolase<br>bypass | WP_010554245 | This study |
| *PpPTA* | *Pseudomonas putida* | Phosphate<br>acetyltransferase | Phosphoketolase<br>bypass | WP_010952014 | This study |
| *CaPDC11* | *Candida albicans* | Pyruvate decarboxylase | Pyruvate bypass | KHC60979 | This study |
| *ScPDC1* | *Saccharomyces*<br>*cerevisiae* | Pyruvate decarboxylase | Pyruvate bypass | NP_013145 | This study |
| *ScPDC5* | *Saccharomyces*<br>*cerevisiae* | Pyruvate decarboxylase | Pyruvate bypass | NP_013235 | This study |
| *ScPDC6* | *Saccharomyces*<br>*cerevisiae* | Pyruvate decarboxylase | Pyruvate bypass | NP_011601 | This study |
| *YIPDC1* | *Yarrowia lipolytica* | Pyruvate decarboxylase | Pyruvate bypass | XP_502647 | (22, 24) |
| *YIPDC2* | *Yarrowia lipolytica* | Pyruvate decarboxylase | Pyruvate bypass | XP_502508 | (22) |
| *ScALD4* | *Saccharomyces*<br>*cerevisiae* | Aldehyde dehydrogenase | Pyruvate bypass | NP_015019 | This study |
| *ScALD6* | *Saccharomyces cerevisiae* | Aldehyde dehydrogenase | Pyruvate bypass | AAB01219 | This study |
| *YIALD1* | *Yarrowia lipolytica* | Aldehyde dehydrogenase | Pyruvate bypass | XP_500380 | (22) |
| *YIALD2* | *Yarrowia lipolytica* | Aldehyde dehydrogenase | Pyruvate bypass | XP_501383 | (22, 24) |
| *YIALD3* | *Yarrowia lipolytica* | Aldehyde dehydrogenase | Pyruvate bypass | XP_503366 | (22) |
| *YIALD4* | *Yarrowia lipolytica* | Aldehyde dehydrogenase | Pyruvate bypass | XP_505802 | (22) |
| *YIALD5* | *Yarrowia lipolytica* | Aldehyde dehydrogenase | Pyruvate bypass | XP_502555 | (22) |
| *YIALD6* | *Yarrowia lipolytica* | Aldehyde dehydrogenase | Pyruvate bypass | XP_504993 | (22) |
| *KIACS1* | *Kluyveromyces lactis* | Acetyl-CoA synthetase | Pyruvate bypass | XP_451146 | This study |
| *PcFACA* | *Penicillium chrysogenum* | Acetyl-CoA synthetase | Pyruvate bypass | XP_056564643 | This study |
| *ScACS2* | *Saccharomyces cerevisiae* | Acetyl-CoA synthetase | Pyruvate bypass | NP_013254 | This study |
| *YIACS1* | *Yarrowia lipolytica* | Acetyl-CoA synthetase | Pyruvate bypass | XP_505057 | (22, 24) |
| *ScCAT2* | *Saccharomyces cerevisiae* | Carnitine acetyltransferase | Carnitine shuttle | CAA78399 | (14, 19) |
| *YICAT2* | *Yarrowia lipolytica* | Carnitine acetyltransferase | Carnitine shuttle | XP_500716 | (21) |
| *YIYAT1* | *Yarrowia lipolytica* | Carnitine acetyltransferase | Carnitine shuttle | XP_505698 | This study |

All genes, except the genes for the carnitine shuttle, were used in the combinatorial screening.

**Fig 4:**
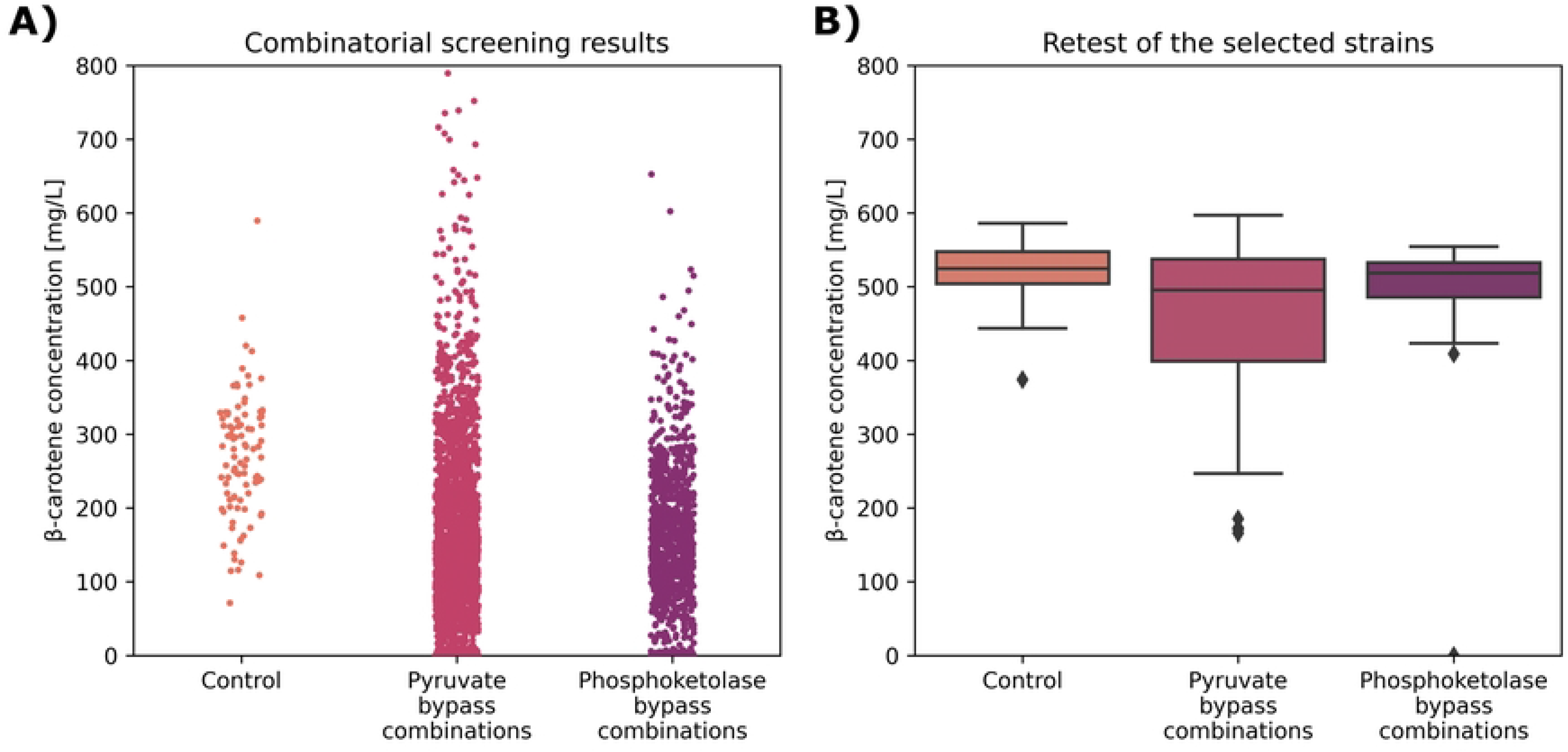
Combinatorial screening results. A) The plot shows beta-carotene production for every transformant from the initial large-scale combinatorial screening in 96-well plates. We selected the top 3% of the strains for retesting (105 strains). B) The results show beta-carotene titers for strains that were selected from the initial screening as the top performers and reassessed in 48-well plates. Both bypasses, pyruvate bypass strains and phosphoketolase bypass strains, performed worse than control strains. Additionally, pyruvate bypass strains showed a larger spread than control and phosphoketolase bypass strains.

**Table 3:**
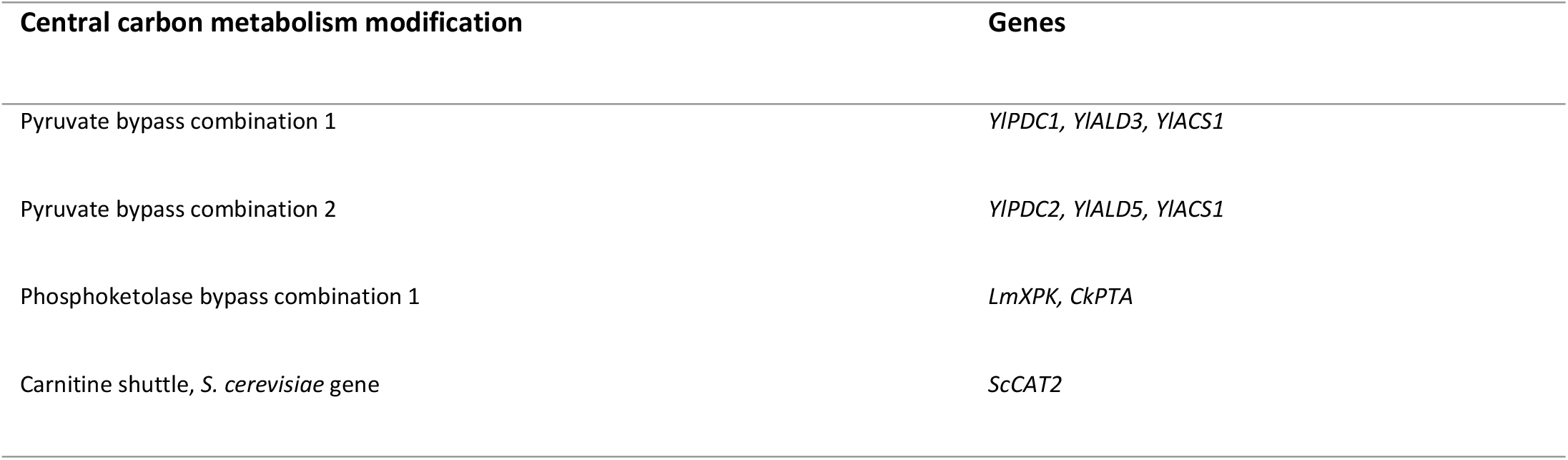
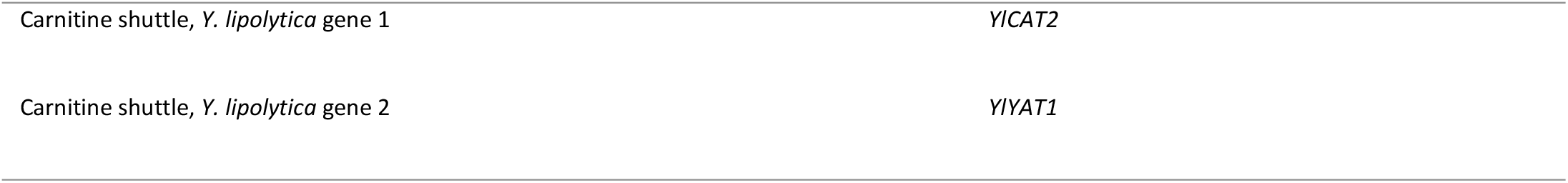
Central carbon metabolism modifications and specific combinations of genes for each of the modifications we tested.

### Selecting a smaller subset of genes for detailed evaluation

We narrowed our focus to six subjects in our study: three genes for the carnitine shuttle (*ScCAT2, YlCAT2,* and *YlYAT1*), two constructs for the pyruvate bypass (*YlPDC1 + YlALD3 + YlACS1* and *YlPDC2 + YlALD5 + YlACS1*), and one construct for the phosphoketolase bypass (*LmXPK + CkPTA*). Firstly, we took two combinations of *Y. lipolytica*’s native genes associated with the pyruvate bypass. Both combinations were previously reported to be beneficial for enhancing the production of polyketides (22). Additionally, we examined a specific combination of genes for the phosphoketolase bypass, which has been reported to improve lipid production (15). Lastly, we investigated three genes for the carnitine shuttle. The first one was the *CAT2* gene from *S. cerevisiae* (14, 19), and two additional ones were *Y. lipolytica*’s native genes that are homologous to the *S. cerevisiae CAT2* gene, namely the *YlCAT2* (21) and *YlYAT1*.

First, we assessed the expression for the chosen subset of genes in the YB-392 strain (Fig 5). For the native genes, expression levels were normalized to that of the actin coding gene *YlACT1* and compared to the native (non-overexpressed) variant. For the heterologous genes, we compared their expression to that of the *YlACT1* gene. Notably, the use of a strong promotor led to elevated expression levels for all genes, particularly for the *YlYAT1* gene, which exhibited expression levels more than 200-fold higher than the native expression level. Once the expression was confirmed for each gene, we proceeded to prepare expression plasmids and introduced them into all the parent strains with the ability to produce beta-carotene (CH-173, TY-160, TY-183, and CH-826). A comprehensive list of the prepared strains can be found in **Error! Reference source not found.**.

**Fig 5:**
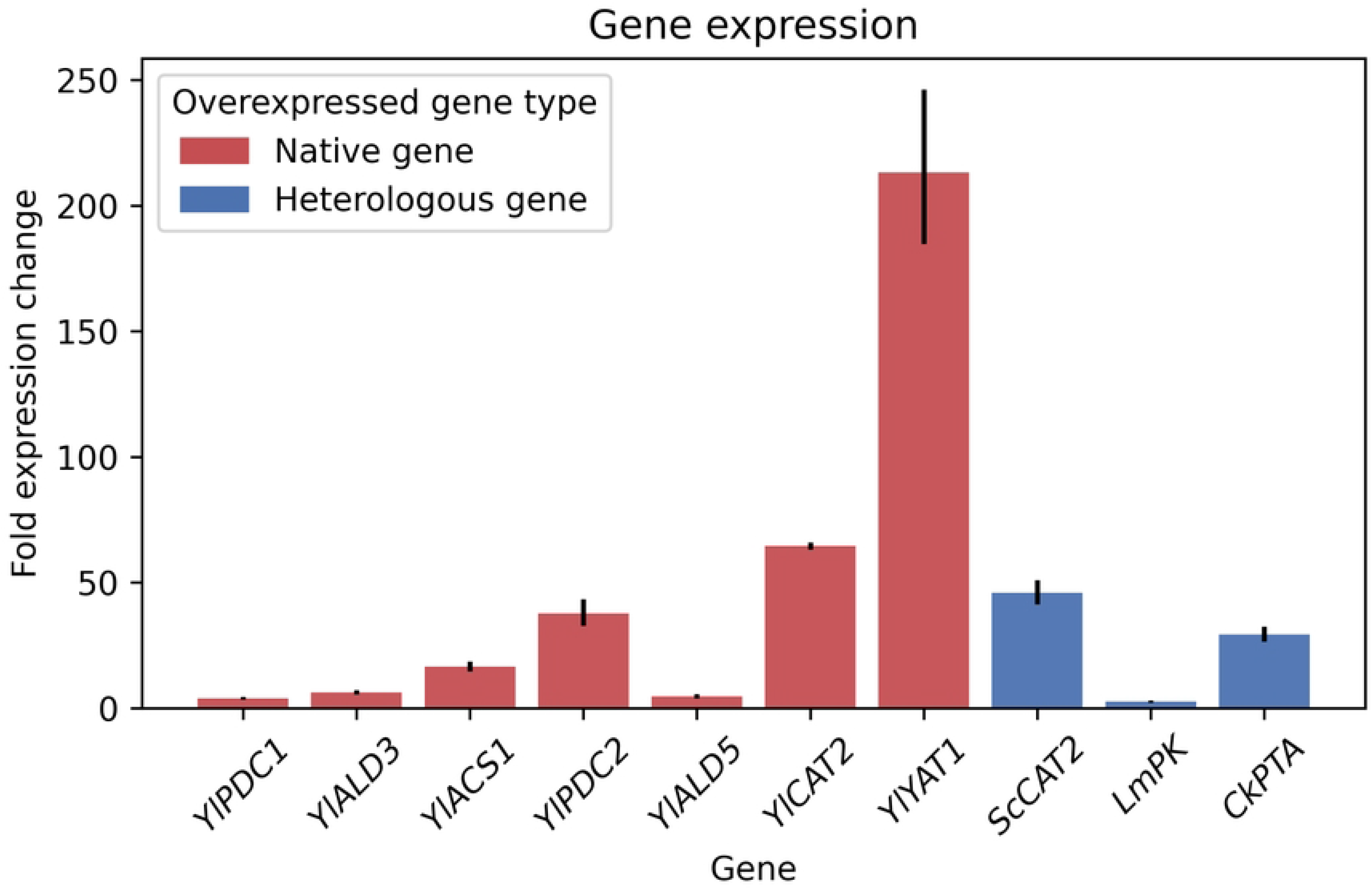
Expression of native and heterologous genes used for central carbon metabolism modifications in the YB-392 strain. The actin coding gene *YlACT1* was used as a reference for normalization, and in the case of heterologous genes also for comparing the expression levels. A functioning expression was confirmed for all genes. Error bars represent standard error, N=3 for each gene.

### Results of the detailed evaluation

All the prepared strains underwent a series of fermentation experiments in shake flasks using defined media. We used one medium with normal glucose concentration and another one with increased glucose concentration to simulate nitrogen limitation. During these experiments, we measured the growth of the strains as well as the concentrations of beta-carotene, acetate, and glucose. To facilitate interpretation of the results, we normalized the data to the control strains. This approach allowed us to gain a broader perspective when assessing the impact of individual modifications on different strains under various conditions.

Our observations revealed that the growth of all strains carrying central carbon metabolism modifications was slightly negatively affected, with the exception of the *YlYAT1* gene transformant, which did not show any growth impairment, or any other change compared to the control strain. For the remaining strains, the growth reduction was marginal, with the growth rates reaching no less than 90% of that observed in the control strain (Fig 6C). In contrast, the production of beta-carotene was significantly impacted by the central carbon metabolism modifications as all the strains containing them exhibited a substantial decrease in their ability to produce beta-carotene. Notably, the pyruvate bypass combination 2 and phosphoketolase bypass showed more than 30% reduced beta-carotene production (Fig 6A). Analyzing the relationship between growth and beta-carotene production, we can conclude that the decline in beta-carotene content (Fig 6D) can mainly be attributed to decreased production rather than growth impairment resulting from the potential metabolic burden imposed by the overexpressed bypasses. Moreover, the yield of beta-carotene (Fig 6B) demonstrated a similar decrease in performance across most of the investigated modifications. However, two modifications, specifically the strains carrying genes for the pyruvate bypass combination 1 or the *YlYAT1* gene, did not show a significant difference in glucose yield for beta-carotene production.

**Fig 6:**
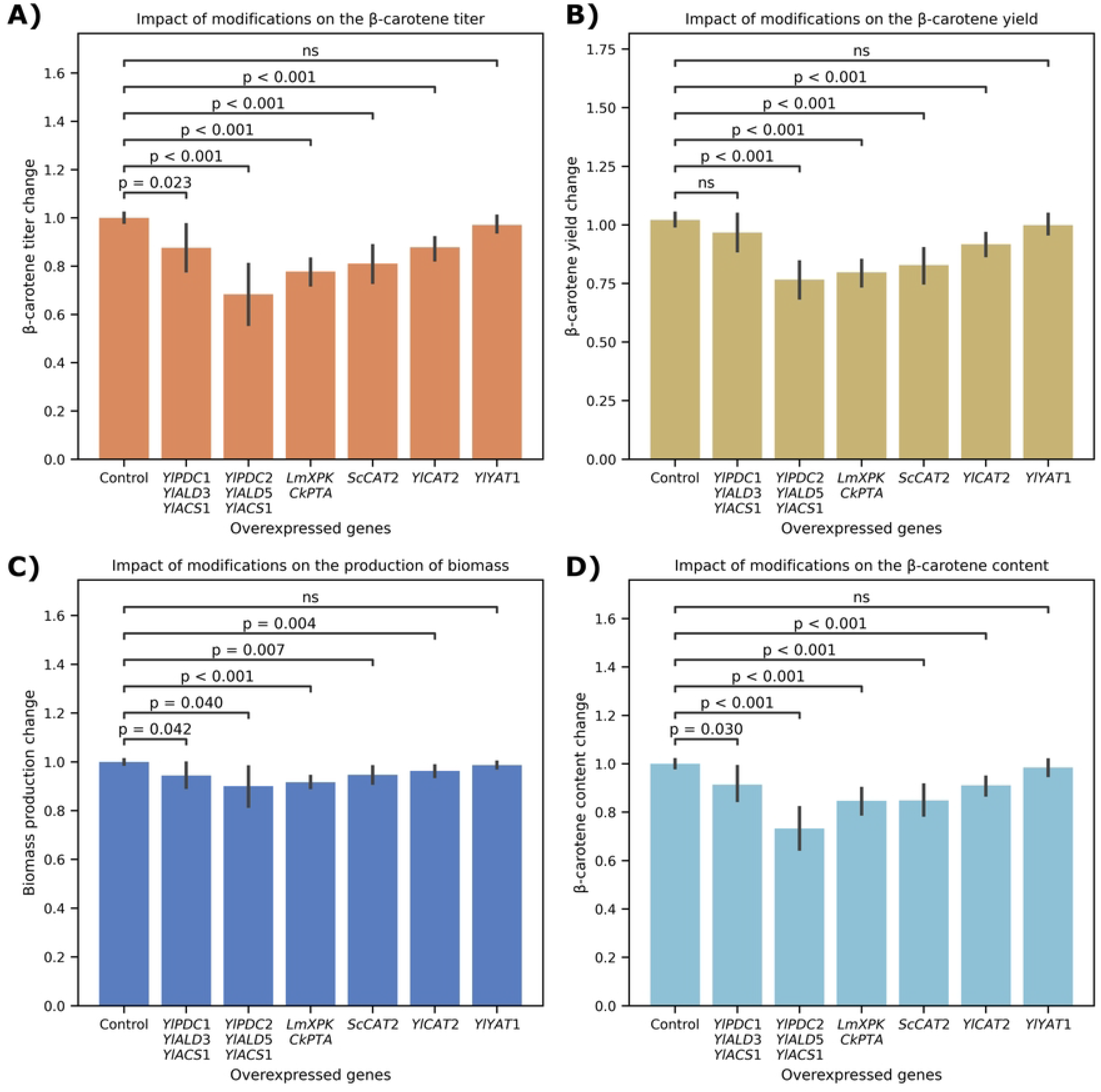
Cumulative results for tested bypasses. Plots above depict normalized cumulative data of multiple experiments (N=3) using multiple sets of strains in two distinct fermentation media (low and high glucose) after 96 hours (results after 72 hours can be found in **Error! Reference source not found.** Fig). In each of the plots, the Control represents parent strains carrying an empty plasmid with just a selection marker gene. Error bars represent confidence intervals with a confidence of 95 %. N=23 for each modification. Plot **A)** focuses on the effect each bypass had on the beta-carotene titer. We can clearly see that neither of the bypasses helped improving the titer. Plot **B)** shows a similar effect on the glucose yield for biosynthesis of beta-carotene. Plot **C)** plot shows that impact of overexpressing each of the bypasses had relatively small impact on the production of biomass, which was approximated by OD600 measurements. Plot **D)** shows the beta-carotene content in Y. lipolytica cells, showing that lowered titer was indeed a consequence of reduced production of beta-carotene and not due to the decreased cell growth. All measurements were taken after a 96 hours of fermentation time.

The observed decrease in yield for the pyruvate bypass combination 2 and the phosphoketolase bypass was unexpected when considering stoichiometry alone, as both pathways are expected to provide a stoichiometrically favorable route towards beta-carotene synthesis. Surprisingly, the pyruvate bypass combination 2, which has been previously identified as the most effective combination of *Y. lipolytica*’s native pyruvate bypass genes for the production of triacetic acid lactone (22), had the most pronounced negative impact on all metrics we investigated in this study. However, Markham and coworkers (22) have suggested that improvements they have observed in the production of triacetic acid lactone may be attributed to a regulatory effect on the redox state of the engineered cells rather than direct effects on the modified acetyl-CoA flux they tried to achieve. These could also imply that the performance improvements seen in other studies investigating the pyruvate bypass in *Y. lipolytica* might be driven by intricate cellular regulatory mechanism rather than solely by the alterations in the carbon flux through the acetyl-CoA.

In contrast, the introduction of phosphoketolase bypass, which has been shown to be largely beneficial for enhancing the production of metabolites downstream of acetyl-CoA in various studies (15–18, 20, 21), led to decreased production, yield, and content of beta-carotene in our study. In three studies that have utilized the same set of genes for the phosphoketolase bypass as we did, the authors have observed positive impact on the production of acetate (17) and lipids (15), and no impact on the production of alpha-humulene (19). However, they have employed different strains of *Y. lipolytica* than we did. These findings emphasize the importance of considering the specific characteristics of each individual *Y. lipolytica* strain, as results obtained from one strain may not be directly applicable to another.

Regarding the modifications involving the carnitine shuttle in *Y. lipolytica*, previous studies done on the W29 derived strains have reported positive effects on the production of lipids (14), beta-farnesene (20), and trans-nerolidol (21). However, Guo and coworkers (19) have reported no effect of *ScCAT2* gene overexpression on alpha-humulene production. In our investigation, we observed a different outcome, revealing a negative impact of overexpressed carnitine shuttle genes on the production, yield, and content of beta-carotene. It is worth noting that the distinct strain background we used (YB-392) may have contributed to the observed differences. However, these results also suggest that the effect of such modifications is highly dependent on the specific end product. The overexpression of *ScCAT2* and *YlCAT2* genes had a similar impact on strain growth, beta-carotene production, yield, and content. Conversely, the strain overexpressing the *YlYAT1* gene showed results most similar to the control strain, with no significant impact on neither growth nor on the beta-carotene production metrics.

Taken together, the above results are largely in agreement with the results we obtained from the initial combinatorial screening where neither of the modifications resulted in the improvement of beta-carotene production, yield, or content. Furthermore, we did not observe an increase in the accumulation of acetate, which could indicate a cellular strategy for regulating acetyl-CoA overflow. We would expect such an overflow to result in increased formation of lipids and consequently in increased beta-carotene production, as such effects have been observed more commonly in *Y. lipolytica* (12).

There are a few considerations we would like to point out. Firstly, it is possible that we were still not able to reach the upper limit of *Y. lipolytica*’s capacity to accommodate carbon flux through acetyl-CoA for production of beta-carotene. Despite our efforts to overcome this limitation by incorporating the CH-826 strain, which has all the major bottlenecks in the pathway towards beta-carotene removed, we did not observe any improvement in the metrics we focused on. Another consideration is the compatibility of genes and the strain background used in our study. It is worth noting that some studies have reported positive effects of similar approaches; however, these studies focused on different end products and utilized different strains and/or genes (Table 1). It is possible that the selection of our specific pool of genes for metabolic bypasses may have resulted in decreased performance. These considerations highlight the complexities and challenges of metabolic engineering in *Y. lipolytica*.

Nonetheless, our findings shed light on the interplay between *Y. lipolytica*’s intrinsic metabolic capabilities and the influence of increased substrate availability. These insights contribute to our understanding of bioprocess optimization and provide valuable knowledge for leveraging *Y. lipolytica*’s natural acetyl-CoA production capacity in industrial applications.

## Conclusion

Our study aimed to assess the impact of metabolic bypasses on the production of beta-carotene in *Y. lipolytica* by modifying the upstream metabolism and increasing the acetyl-CoA supply. However, the modifications we investigated in central carbon metabolism did not result in improved beta-carotene production. Surprisingly, they mostly had a negative impact on its production. Our findings, along with previous research, suggest that the bottlenecks in the production of carotenoids with oleaginous yeast *Y. lipolytica* are likely to be located elsewhere and that factors such as strain background, genetic compatibility, regulatory mechanisms, and especially the choice of the end product can greatly influence the outcomes of metabolic pathway engineering.

## Materials and methods

### Strains, materials, media

Plasmid construction and propagation were performed using the *Escherichia coli* DH10β strain. For LB medium 10 g peptone, 5 g yeast extract, and 5 g NaCl was dissolved in 1000 mL dH_2_O and autoclaved. After sterilization, the appropriate antibiotic was added for plasmid selection.

All *Y. lipolytica* strains used for this study originated from the YB-392 wild-type strain (11) and are listed in **Error! Reference source not found.**. For the carbon source, a 60 % (w/v) stock glucose solution was prepared by dissolving 600 g/L glucose in dH2O. Sterilization was achieved by autoclaving. The stock solution was stored at 4 °C. A 50x stock MES buffer solution was prepared by dissolving 97.6 g 2-ethanesulfonic acid in 500 mL dH_2_O and adjusting pH to 5.7 before filtration through a 0.2-micron filter. The stock solution was stored at 4 °C. YPD medium was prepared by dissolving 20 g peptone and 10 g yeast extract were dissolved in 800 mL dH_2_O, adjusting pH to 6.5, and autoclaving. After sterilization, 33.3 or 166.7 mL of the 60 % stock glucose solution was added to achieve a final glucose concentration of 20 or 100 g/L (YPD or YPD100, respectively). Mineral media was prepared by dissolving 1.7 g yeast nitrogen base without amino acids and without ammonium sulphate (SIGMA) and 0.4 g (NH_4_)_2_SO_4_ in 800 mL dH_2_O, followed by autoclaving. After sterilization, 33.3 mL or 100 mL of the 60 % stock glucose solution (SC-2 or SC-5, respectively), 20 mL of the 50x stock MES buffer solution, and sterile dH_2_O were added to adjust the total volume to 1.0 L. Yeast Synthetic Drop-out medium supplements (SIGMA) were prepared according to manufacturer’s recommendations. Selection with dominant markers was performed with addition of 300 µg/mL nourseothricin or 400 µg/mL hygromycin. Solid plates were prepared by adding 20 g/L agar.

### Cloning and transformation

For the construction of expression cassettes, the YTK system adapted from (25) was employed. All transcription units containing the genes of interest were assembled with a strong constitutive P_hp8d_ promoter and a T_cyc1_ terminator. The final expression cassettes were generated by combining the appropriate transcription units with zeta regions and the selection marker. The heterologous genes used in this study were codon optimized for transcription in *Y. lipolytica* and obtained from Twist Bioscience. The genes were verified by sequencing, and a list of the genes is provided in Table 2 and Table 4. Similarly, a list of all assembled expression cassettes can be found in **Error! Reference source not found.**. Before transformation, the expression cassettes were linearized through restriction digestion and the linearized fragments of correct length were purified using gel electrophoresis.

**Table 4:**
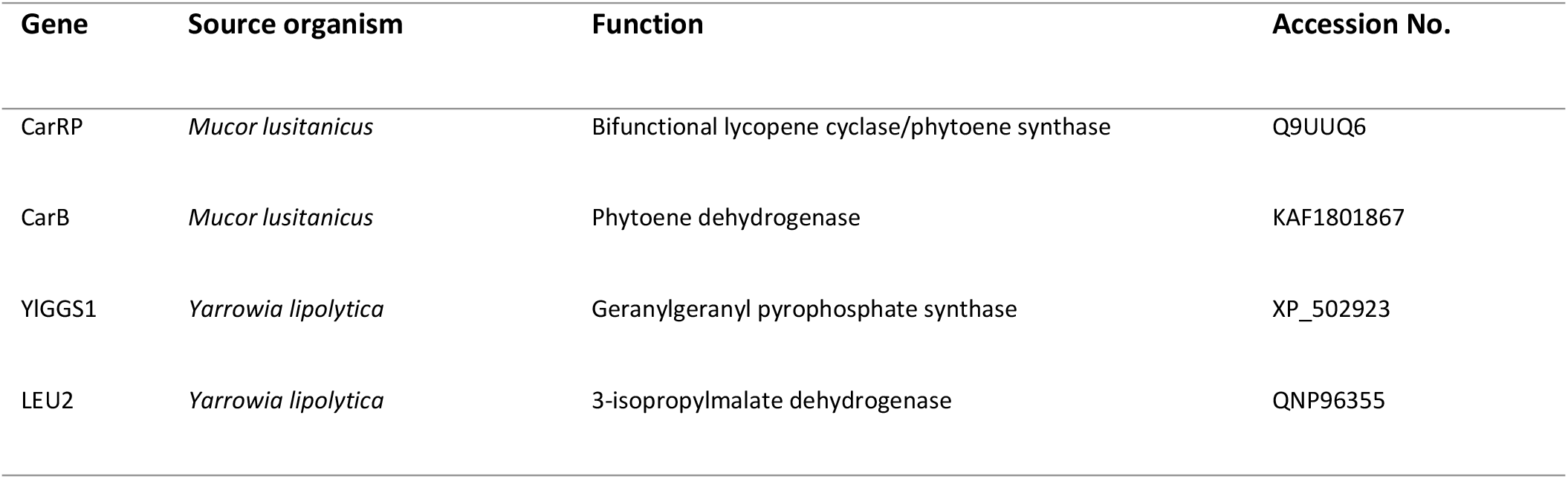
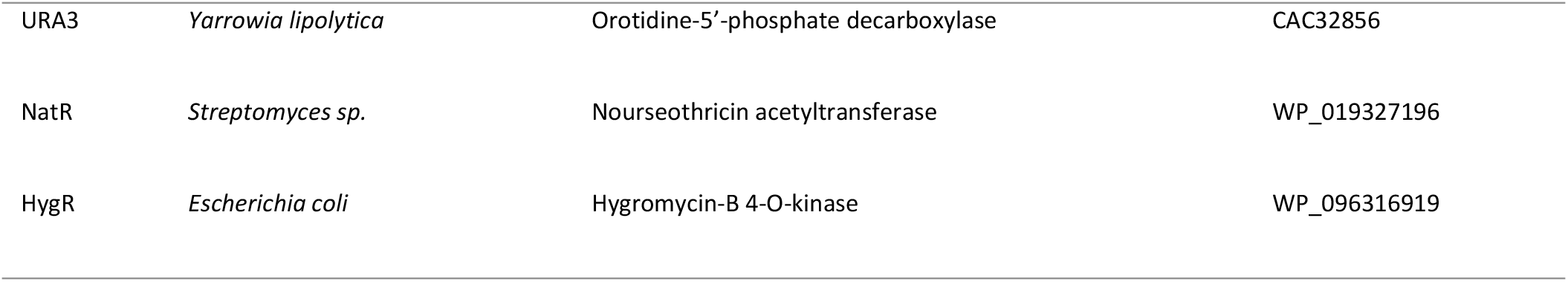
Other genes used in this study accompanied with their source organism, function, and accession number.

| Gene | Source organism | Function | Accession No. |
| --- | --- | --- | --- |
| CarRP | *Mucor lusitanicus* | Bifunctional lycopene cyclase/phytoene synthase | Q9UUQ6 |
| CarB | *Mucor lusitanicus* | Phytoene dehydrogenase | KAF1801867 |
| YIGGS1 | *Yarrowia lipolytica* | Geranylgeranyl pyrophosphate synthase | XP_502923 |
| LEU2 | *Yarrowia lipolytica* | 3-isopropylmalate dehydrogenase | QNP96355 |
| URA3 | *Yarrowia lipolytica* | Orotidine-5'-phosphate decarboxylase | CAC32856 |
| NatR | *Streptomyces sp.* | Nourseothricin acetyltransferase | WP_019327196 |
| HygR | *Escherichia coli* | Hygromycin-B 4-O-kinase | WP_096316919 |

*Y. lipolytica* strains were transformed with the linearized fragments using the adapted LiAc transformation protocol (26). Positive transformants were selected on appropriate medium and checked for correct gene integration through colony PCR. To assess expression levels, RNA was extracted from 24-hour-old cultures using the Qiagen RNA extraction kit and following manufacturer’s instructions. Reverse transcription and genomic DNA digestion were performed using the Thermo Scientific Reverse Transcription kit. The qPCR analysis was carried out using the Applied Biosystems qPCR instrument with the Thermo Scientific Maxima SYBR Green/ROX qPCR Master Mix. The gene encoding actin (*YlACT1*) served as the normalization control. All PCRs were performed in triplicate. The primers used for colony PCR and qPCR can be found in **Error! Reference source not found.**.

## 96-well deepwell plate cultivation for the combinatorial screening

For the first combinatorial screening we prepared a pre-culture by incubating 500 µL of YPD medium in each well of a 96-well microtiter plate for 24 hours on a rotary shaker set at 28 °C, 220 rpm, and 70 % humidity control. After 24 hours, 10 µL of seed culture was transferred in 500 µL of fresh YPD100 in the 96-well microtiter plate. The fermentation was carried out for four days on a rotary shaker set at 28 °C, 220 rpm, and 70 % humidity control. After 96 hours, samples were taken for measurements.

## 48-well deepwell plate cultivation for re-evaluation of selected strains from the combinatorial screening

For the screening of selected strain from combinatorial screening we prepared a pre-culture by incubating 1000 µL of YPD medium in each well of a 48-well microtiter plate for 24 hours on a rotary shaker set at 28 °C, 220 rpm, and 70 % humidity control. After 24 hours, 20 µL of seed culture was transferred in 1000 µL of fresh YPD100 in the 48-well microtiter plate. The fermentation was carried out for four days on a rotary shaker set at 28 °C, 220 rpm, and 70 % humidity control. After 96 hours, samples were taken for measurements.

## 48-well deepwell plate cultivation for assaying beta-carotene production by parent strains

For assaying the beta-carotene production by parent strains, we prepared a pre-culture by incubating 1000 µL of YPD medium in each well of a 48-well microtiter plate for 24 hours on a rotary shaker set at 28 °C, 220 rpm, and 70 % humidity control. After 24 hours, 20 µL of seed culture was transferred in 1000 µL of fresh SC-5 in the 48-well microtiter plate. The fermentation was carried out for four days on a rotary shaker set at 28 °C, 220 rpm, and 70 % humidity control. After 96 hours, samples were taken for measurements.

### Shake-flask fermentation

A pre-culture was prepared by incubating 5 mL of YPD medium for 24 hours on a rotary shaker set at 28 °C, 220 rpm, and 70 % humidity control. After 24 hours, cell density was measured at 600 nm using a Tecan Infinite M200 plate reader instrument. For the main fermentation, 250 mL Erlenmeyer flasks containing 50 mL of mineral medium were inoculated with pre-cultures to achieve an initial optical density (OD_600_) of 0.05. The shake-flask fermentation was carried out for four days on a rotary shaker set at 28 °C, 220 rpm, and 70 % humidity control. After 72 and 96 hours, samples were taken for measurements.

### Carotenoid extraction and quantification

For the extraction of beta-carotene, a 400 µL sample of the fermented broth was mixed with an equal volume of butyl acetate and 200 µL of glass beads. The mixture was then homogenized using a FastPrep-96 instrument at medium speed for 5 minutes. After homogenization, the samples were centrifuged at 14000 rpm for 10 minutes, and the resulting supernatant was carefully transferred to glass vials for further analysis using HPLC.

Quantification of beta-carotene was performed using the Thermo Scientific Accela HPLC machine. Separation was performed on the Kromasil 100-5 C18 column, 50 x 4.6 mm, 5 µm particle size. The mobile phase program started with 20% dichloromethane and 80% methanol, with linear gradient increase to 50% of dichloromethane in 10 minutes. The flow rate was set at 0.5 mL/min, and the column oven temperature was maintained at 25 °C. Detection of beta-carotene was achieved using a DAD detector, set at wavelength of 455 nm.

## Acknowledgments

We would like to express our gratitude to all individuals who contributed, directly or indirectly, to the successful execution of this study.

